# Despite reproductive interference, the net outcome of reproductive interactions among spider mite species is not necessarily costly

**DOI:** 10.1101/113274

**Authors:** Salomé H. Clemente, Inês Santos, Rita Ponce, Leonor R. Rodrigues, Susana A. M. Varela, Sara Magalhães

## Abstract

This preprint has been reviewed and recommended by Peer Community in Evolutionary Biology (http://dx.doi.org/10.24072/pci.evolbiol.100025). Reproductive interference is considered a strong ecological force, potentially leading to species exclusion. This supposes that the net effect of reproductive interactions is strongly negative for one of the species involved. Testing this requires a comprehensive analysis of interspecific reproductive interactions, accounting for the order and timing of mating events, and for their effects on either fertility or fecundity. To this aim, we measured reproductive interactions between a focal species, *Tetranychus urticae*, and an invasive (*T.evansi*) and a resident (*T. ludeni*) species, varying the mating sequence and interval, and measuring the effect of such crosses on fecundity and offspring sex ratio (a measure of fertility, as these species are haplodiploid). We found that mating with heterospecifics affected fecundity and sex ratio negatively, but also positively, depending on the species involved, and on the order and timing of mating events. Overall, the net effect of reproductive interactions was weak despite strong effects of particular events. In natural situations, the outcome of reproductive interactions will thus hinge upon the frequency of each event.

## Introduction

Reproductive interference, that is, any kind of sexual interaction between two species that diminishes the fitness of at least one of them (Gröning and Hochkirch 2008, Kishi et al. 2009, Burdfield-Steel and Shuker 2011), may have severe effects on the outcome of species interactions. Indeed, theory predicts that reproductive interference may contribute to species exclusion more often than resource competition (Gröning and Hochkirch 2008, Kishi et al. 2009, Kishi and Nakazawa 2013). For example, it has been posited that reproductive interference may underlie the success of some invasive species (e.g. Nishida et al. 2012).

Most studies of reproductive interference concern the fitness outcome of interspecific matings of two species that do not produce viable hybrids (Gröning and Hochkirch 2008). In this case, the reproductive effects of the interspecific interaction will be expressed only when organisms mate with both conspecifics and heterospecifics (as mating with heterospecifics alone will yield no offspring). Moreover, clearly evaluating the effects of reproductive interference on exclusion in polyandrous species necessitates measuring all possible combinations of mating order (*i.e*., whether heterospecific matings occur before or after conspecific ones) and timing (*i.e*., the interval between mating events) between pairs of species. It is also important to test whether reproductive interactions affect fecundity (egg production) or fertility (egg fertilization). This information can then be integrated to predict the net outcome of reproductive interactions between species. Despite the many studies on reproductive interference, none has yet applied this approach. Indeed, some studies attempt to predict how reproductive interference affects species exclusion, but do so by focussing on some sequence events only. For example, Takafuji (1997) used a Lotka-Volterra modified model to predict the effect of reproductive interference between two mite species (*Panonychus citri* and *P. mori*) on species exclusion, but they used only one possible combination of mating interactions between species. In contrast, other studies consider different orders of mating events (eg, Kyogoku and Nishida 2013), but do not integrate this information to generate a prediction concerning the net effect of reproductive interactions on species distributions.

Here, we aimed at testing how the outcome of different mating events among species may affect their life-history traits, using spider mites, a group where reproductive interference has been frequently observed (Collins and Margolies 1994; Takafuji et al. 1997; Ben-David et al. 2009, Sato et al. 2014). Spider mites are haplodiploid, hence the distinction between fecundity and fertilization effects can be made given that fertilized eggs result in female offspring and unfertilized eggs in male offspring. Thus, fertilization failures can be detected by a reduction in the proportion of female offspring, whereas impairment of egg production is detected by a reduction in the total number of offspring.

We used a system composed of one focal species, *Tetranychus urticae*, in sexual heterospecific interactions with another resident species, *T. ludeni*, and an invasive species, *T. evansi*. These three herbivorous species co-occur in the Mediterranean region and are often found on the same host plant (Escudero and Ferragut 2005, Boubou et al. 2012, Godinho et al. 2016). *T. urticae* and *T. ludeni* are resident species, and *T. evansi* has recently invaded the European continent, via Portugal (Boubou et al. 2012). Heterospecific matings among these species have been observed, although they mostly do not result in viable offspring (Sato et al. 2014, 2016, Clemente et al. 2016, S. Magalhães pers. obs.). Moreover, *T. evansi* can exclude *T. urticae* on tomato plants (Sarmento et al. 2011a), a result that correlates with field observations (Ferragut et al. 2013). Finally, a recent study has shown that, in competition with *T. evansi*, the population growth of *T. urticae* is more severely affected when plants are colonized by virgin females than when plants are colonized by mated females, suggesting that reproductive interference may be responsible for the species distribution patterns observed (Sato et al. 2014).

To postulate hypotheses concerning the consequences of heterospecific matings, it is crucial to understand within-species reproductive behaviour. *T. urticae*, the focal species, exhibits first male sperm precedence, with second matings being sometimes effective if they occur within the 24 hours following the first (Helle 1967). However, females that mate multiple times with conspecific males, after a 24h interval between matings, produce fewer fertilized offspring (*i.e*., females) (Macke et al. 2012), suggesting that sperm displacement after 24h is possible. Here, we hypothesize that mating order and the mating interval will affect the outcome of reproductive interference in *T. urticae*. Also, given that *T. evansi*, the invasive species, displaces *T. urticae*, unlike *T. ludeni*, we expect the former to exert stronger effects than the latter. To this aim, we performed crosses between *T. urticae* and the two other species at different time intervals and with different mating orders, and measured the consequences for the two species involved in the cross.

## Material and Methods

### Stock Cultures

The mite species used in this study were collected in Carregado (39.022260, - 8.966566), Portugal, and all laboratory populations were established from an initial pool of 300 mated females. The laboratory population of *T. urticae* was collected on tomato plants *(Solanum lycopersicum)* in May 2010, that of *T. evansi* on *Physalis angulata* in May 2012 and that of *T. ludeni* on tomato in September 2012. The populations of *T. evansi* and *T. ludeni* became extinct in August 2012 and May 2013, respectively, being subsequently replaced with populations from the same location, both collected in *Datura stramonium* plants. Both populations of *T. evansi* and *T. ludeni* were used in the experiments.

Species identity was confirmed through polymerase chain reaction-restriction fragment length polymorphism (PCR-RFLP) of the ITS2 region (Hurtado et al. 2008), on approximately 50 females of each population. Total genomic DNA was extracted from each individual spider mite using the Sigma-Aldrich GenEluteTM Mammalian Genomic DNA Miniprep Kit, following manufacturer’s instructions, except for the elution volume, which we set to 20μL of RNase free water (Qiagen NV, Venlo, The Netherlands) to increase the concentration of DNA obtained from this very small animal (c.a. 300μm long).

Adult females from populations used in this experiment were screened for *Wolbachia* using the primers *wsp (Wolbachia-specific* primers) 81F and 691R (Braig et al. 1998). We did this to avoid potential cytoplasmic incompatibility as a confounding factor in our measurements. PCR assay procedures were as described in Breeuwer (1997). Results were positive for *Wolbachia* infection and all spider mite populations were thus treated by placing adult females in detached bean leaves with tetracycline (0.025% w/v) for three consecutive generations, then absence of *Wolbachia* was confirmed using the same protocol as above. Other endosymbionts tested (Arsenophorous, Rickettsia, Spiroplasma and Cardinium) were absent from these populations.

Bean *(Phaseolus vulgaris)* and tomato *(Solanum lycopersicum)* plants were planted every week and grown in an herbivore-free greenhouse, being watered two to three times a week. *T. urticae* populations were maintained on trays with 6-10 bean plants whereas those of *T. evansi* and *T. ludeni* were kept on tomato plants at 25°C, both with a 16 L: 8D photoperiod. Plant trays were changed every two weeks, placing old leaves on top of uninfested plants. Cultures were kept inside plastic boxes (28x39x28 cm), with an opening of 25x15 cm polyamide fabric (80 μm mesh width).

### Experimental procedure

Experiments were done on the plant species from which the female tested had been cultured. As in the literature there was no information on whether hybridization is possible between *T. urticae* and *T. ludeni*, we studied the outcome of a single heterospecific mating between these two species (the same analysis for *T. urticae* and *T. evansi* was performed in a previous experiment (Clemente et al. 2016)). Subsequently, we set out to study the heterospecific interactions between *T. urticae* and the invasive *T. evansi* and the resident *T. ludeni* species for which we analysed the outcome of mating with a heterospecific male before or after a conspecific male. Since we focused on interactions with *T. urticae* (the focal species of our study), we performed crosses between *T. urticae* males or females and *T. evansi* or *T. ludeni* males or females, but not between the two latter species. All experiments were performed in an acclimatized room at approximately 25°C.

#### a) The outcome of a single heterospecific mating between *T. urticae* and *T. ludeni*

To determine whether hybridization occurred between *T. urticae* and *T. ludeni*, we measured the offspring sex-ratio resulting from single heterospecific matings. Given that only females develop from fertilized eggs, a whole-male offspring would mean unsuccessful hybridization. However, even in the absence of viable hybrids, heterospecific matings could result in aborted development of heterospecifically-fertilized eggs, meaning that females would produce fewer eggs. To test this, we compared the fecundity of *T. urticae* and *T. ludeni* females that mated with a heterospecific male to that of virgin females and of females mated with a conspecific male.

Females were collected from the stock populations, isolated at the quiescent deutonymph stage (which precedes their last moult before reaching adulthood), and kept in groups of approximately 15 females on bean (*Phaseolus vulgaris*) leaf discs (2 cm^2^) until emergence, to ensure their virginity. Adult males were collected from the same stock populations and kept isolated in leaf discs (2 cm^2^) for at least 24 hours before the assay, to ensure sperm replenishment. Females were placed individually in leaf discs (1 cm^2^) with either a conspecific or a heterospecific male and observed continuously until copulation occurred. Only matings that lasted at least 1 minute were considered effective (Boudreaux 1963). These experiments had the maximum duration of 2 hours. If no mating occurred within this time, individuals were discarded. Subsequently, females were isolated in a leaf disc (2 cm^2^), then transferred to a new disc every three days until the female’s death. The number of eggs laid was registered after female transfer to a new leaf disc. Eggs were left to develop until adulthood when offspring sex-ratio could be determined. With this data, we tested whether heterospecific matings affected (a) the mean daily fecundity and (b) offspring sex ratio (hence the proportion of fertilized offspring).

#### b) The outcome of heterospecific matings that precede or follow conspecific ones

To determine the outcome of mating with a heterospecific male before or after a conspecific male between *T. urticae* and the other two species, we compared the fecundity and offspring sex ratio of those crosses to that of females that mated with two conspecific males. The experimental procedure was as described above, except that we let females mate with a conspecific or a heterospecific male, then placed the focal females with another male. We created the following mating sequences: conspecific-conspecific, conspecific-heterospecific and heterospecific-conspecific. The second mating occurred either immediately after the first mating (0 hours treatment) or 24 hours later. If no mating was observed within 2 hours, the females were discarded. We used the 0h and 24h mating intervals because the time interval was shown to affect the degree of sperm precedence in spider mites (Helle 1967).

### Statistical analysis

All analyses were carried out using R (version 3.3.2, R Development Core Team 2016). To analyse female fecundity within each species (*T. urticae*, *T. evansi* and *T. ludeni*), we used linear models (LM procedure), considering the mean number of eggs per day as the response variable (oviposition rate). To analyse offspring sex ratio within each species, we used generalized linear models (GLM procedure) with a quasi-binomial distribution - due to overdispersion of the data -, considering the number of female and male offspring produced by each focal female as the response variables (analysed together with the function cbind).

For both types of analyses, we used as fixed factors the mating order (with three levels: the control treatment, where a female mated twice with conspecific males; an experimental treatment where the heterospecific male was the first to mate with the female; and another experimental treatment where the heterospecific male was the second to mate with the female) and the mating interval (with two levels: either 0h or 24h interval between matings). We also tested the interaction among these fixed factors. If the interaction was non-significant, a backward stepwise procedure was used to find the best simplified fitted model. We performed independent analyses for each species within each species pair (i.e. for *T. urticae* and *T. evansi* females in *T. urticae versus T. evansi* crosses; and for *T. urticae* and *T. ludeni* females in *T. urticae versus T. ludeni* crosses).

We did a first block of experiments with the populations of *T. evansi* and *T. ludeni* collected in 2012 (block 1). For question b) we also did a second block of experiments with populations of those species from 2013 (block 2). In block 2 we did not repeat all treatments, but only the crosses that were not complete before the extinction of block 1 populations, as well as their respective controls – hence, there were no treatments that were only performed in block 2. Because of that, instead of including the factor block in the statistical models as a covariate, we did all the statistical analyses with block 1 only and with block 1 and block 2 together. Since the results were qualitatively similar (Table S1), here we present the results from the analysis with block 1 and block 2 together.

## Results

### a) The outcome of a single heterospecific mating between *T. urticae* and *T. ludeni*

Crosses between *T. ludeni* and *T. urticae* resulted in 100% male offspring, indicating that hybrid production between these species is inexistent. The fecundity of *T. urticae* females that mated heterospecifically was not significantly different from that of virgin females or from that of females mated with a conspecific male (F_2,78_= 1.886, P= 0.1585; Figure 1). On the other hand, the fecundity of *T. ludeni* females that mated with conspecifics or heterospecifics was significantly higher than that of virgin females (F_2,66_= 5.636, P= 0.0055; Figure 1). Therefore, mating with heterospecific males does not result in the aborted fertilization of oocytes for *T. urticae* and *T. ludeni* females.

**Figure 1.**
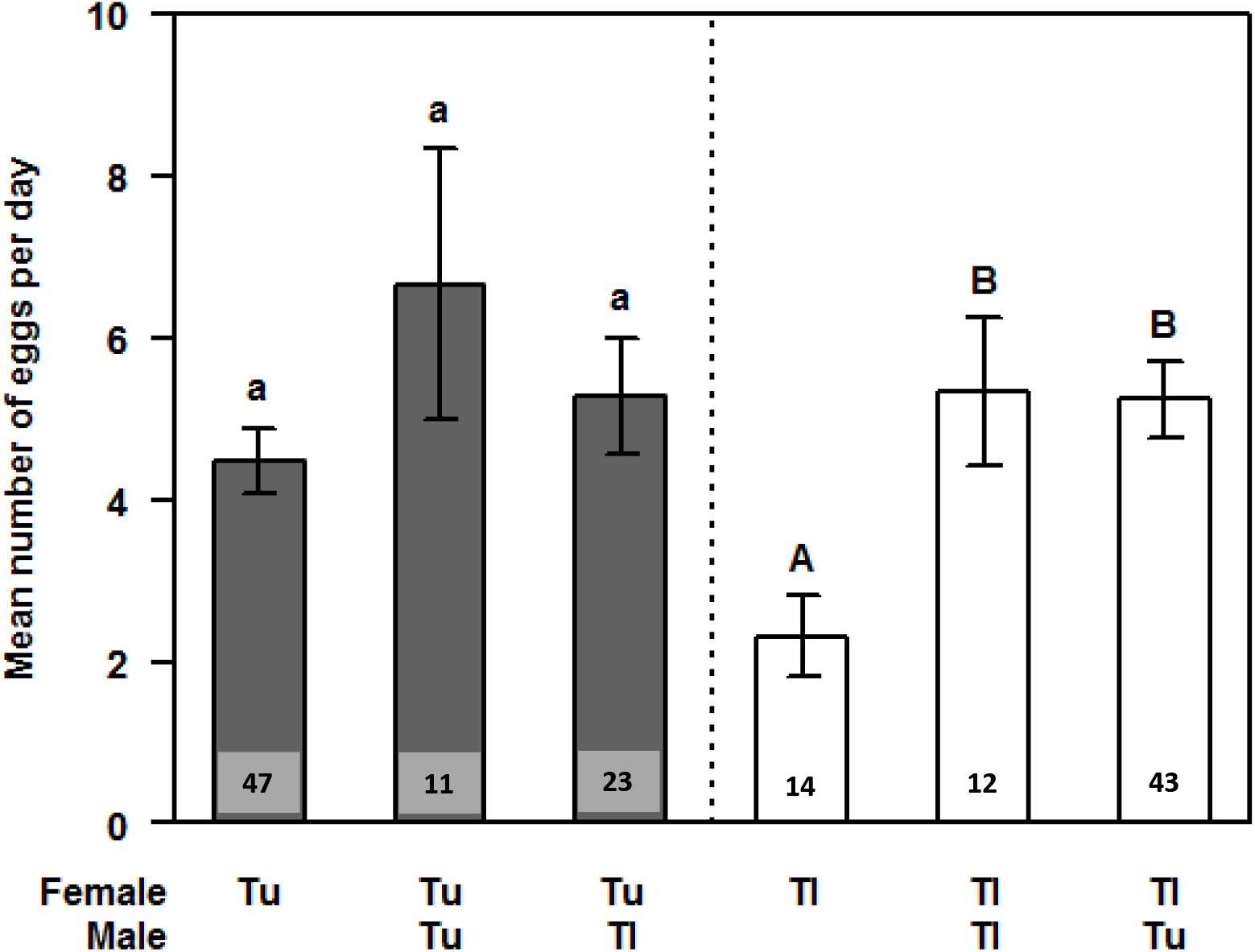
Average daily fecundity of virgin females, and of females that have mated with a conspecific or a heterospecific male. Tu: *T. urticae* males or females; Tl: *T. ludeni* males or females. Grey bars: matings involving *T. urticae* females; white bars: matings involving *T. ludeni* females. Error bars represent the standard errors of the mean. Numbers on the bottom of bars represent the sample size for each type of mating.

### b) The outcome of heterospecific matings that precede or follow conspecific ones

#### (i) T. urticae vs T. evansi

The oviposition rate of *T. urticae* females that mated with either a conspecific and a heterospecific or with two conspecific mates varied significantly according to mating order in interaction with mating interval (F_2,136_ = 6.026, P = 0.0031). Specifically, it was higher for *T. urticae* females that mated with *T. evansi* males just before mating with a conspecific male than for any other cross at 0h mating interval (|t|(1) = 4.964,P < 0.0001 and |t|(1) = 3.288, P = 0.0009, in comparison with double conspecific matings and with matings with a conspecific followed by a mating with an heterospecific, respectively; Fig. 2a). At the 24h interval, however, mating combinations did not affect this trait. The proportion of fertilized offspring (*i.e*., daughters) of females *T. urticae* also varied significantly according to mating order in interaction with mating interval (F_2,106_= 57.219, P= 0.007). But in contrast to the oviposition rate, this trait was affected at the 24h interval only, in which mating with a *T. evansi* male after mating with a conspecific male resulted in a decrease in the proportion of fertilized offspring of *T. urticae* females, relative to other mating sequences (|t|(1) = 5.362, P < 0.0001 and |t|(1) = 5.103, P < 0.0001, in comparison with double conspecific matings and with matings with an heterospecific followed by a mating with a conspecific male, respectively; Fig. 2b).

**Figure 2.**
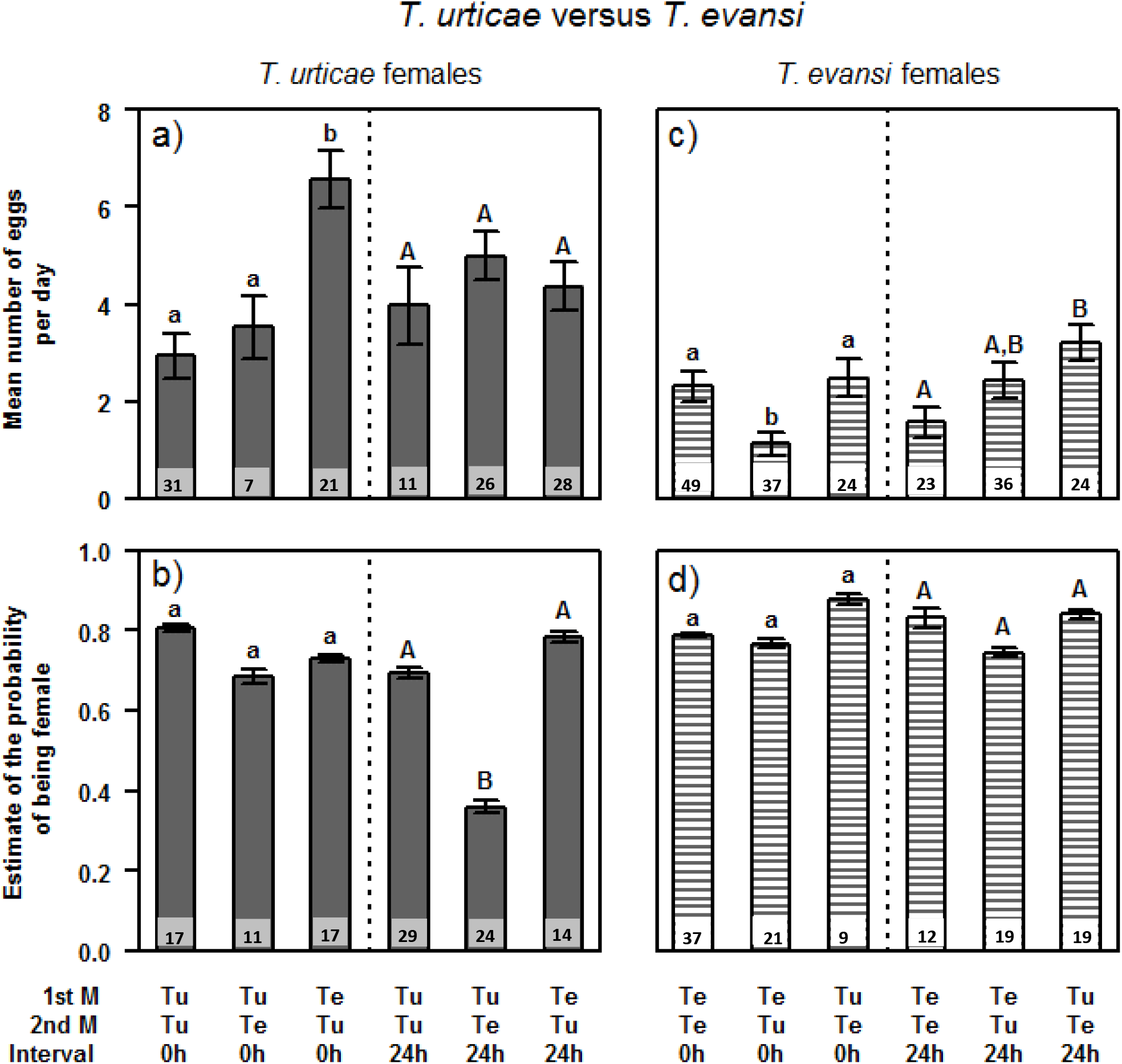
Average daily fecundity and estimated offspring sex ratio resulting from interactions between *T. urticae* (a, b; grey solid bars) and *T. evansi* (c,d; striped bars) females with conspecific and heterospecific males. In each plot, bars on the left side of the dotted straight line correspond to treatments where second matings occurred immediately (0h) after the first one; bars on the right side correspond to treatments where second matings occurred 24h after the first one. “1st M”: first male that mated with the female; “2nd M”: second male. The interval indicates the time of occurrence of the second mating, i.e., if immediately after the first mating (0h) or 24h later. “Tu”: *T. urticae* males; “Te”: *T. evansi* males. Letters above the bars indicate significant differences among treatments (small letters: among crosses occurring with a 0h interval; capital letters: among crosses occurring with a 24h interval). Error bars represent the standard errors of the mean. For offspring sex ratio, we obtained the estimates of the probability of being female and correspondent standard errors of the mean from the statistical GLM models. This takes into account sex ratio variation among females, as well as the quasi-binomial correction for overdispersion of the data. Numbers on the bottom of bars represent the sample size for each type of mating.

The mating order also affected differentially the oviposition rate of *T. evansi* females, depending on the interval between matings (F_2,187_= 4.977, P= 0.0078). *T. evansi* females that mated with *T. urticae* males immediately after conspecific mates had reduced oviposition rate relative to other mating sequences at this time interval (|t|(1) = 2.841, P = 0.0050 and |t|(1) = 2.692, P = 0.0078 in comparison with double conspecific matings and with matings with a heterospecific followed by a mating with a conspecific male, respectively; Fig. 2c); however, if the heterospecific cross occurred 24 hours before the conspecific cross, the oviposition rate of *T. evansi* females increased relative to double conspecific matings at this time interval (|t|(1) = 2.948, P = 0.0036; Fig. 2c). These crosses did not significantly affect sex ratio (F_2,111_=3.786, P=0.6923; Fig. 2d).

#### (ii) T. urticae vs T. ludeni

In crosses with the resident species (*T. ludeni*), the oviposition rate of *T. urticae* females varied significantly according to mating order in interaction with mating interval (F_2,144_ = 3.694, P = 0.0273). Specifically, we found that, at 0h interval, females that mated first with a conspecific then with a heterospecific male had higher oviposition rate than females that mated first with a heterospecific then with a conspecific male (|t|(1) = 2.736, P = 0.0070; Fig. 3a). At the 24h interval, the oviposition rate of females that mated first with a conspecific then with a heterospecific male was lower than that of double conspecific crosses. (|t|(1) = 2.505, P = 0.0134; Fig. 3a). *T. urticae* females suffered no significant changes in offspring sex ratio from matings with *T. ludeni* males (F_2,99_ = 10.769, P = 0.3195; Figure 3b).

**Figure 3.**
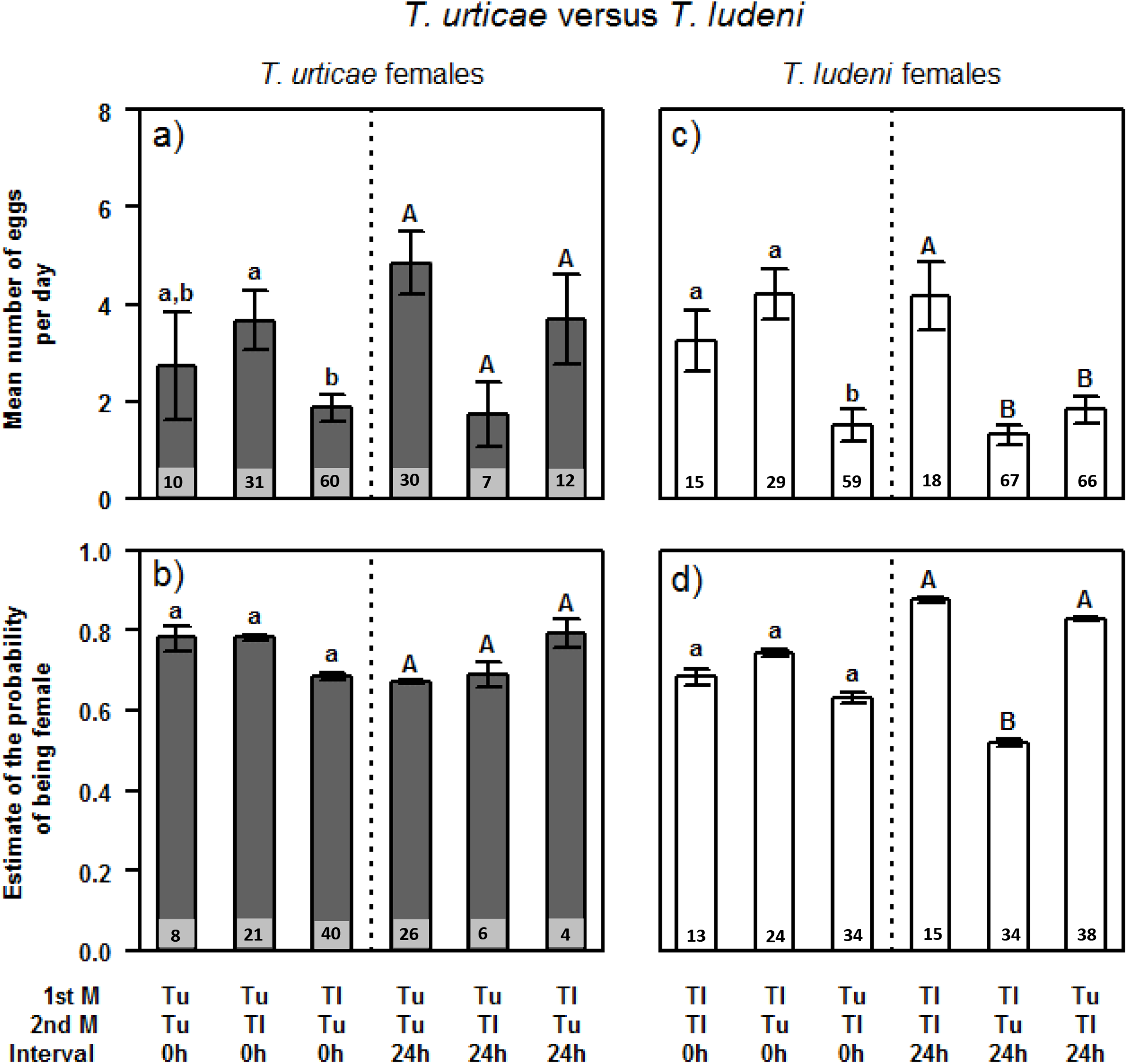
Average daily fecundity and estimated offspring sex ratio resulting from interactions between *T. urticae* (plots a, b; grey bars) and *T. ludeni* (plots c, d; white bars) females with conspecific and heterospecific males. In each plot, bars on the left side of the dotted line correspond to treatments where second matings occurred immediately (0h) after the first one; bars on the right side correspond to treatments where second matings occurred 24h after the first one. “1st M”: first male that mated with the female; “2nd M”: second male. The interval indicates the time of occurrence of the second mating, i.e., if immediately after the first mating (0h) or 24h later. “Tu”: *T. urticae* males; “Tl”: *T. ludeni* males. Letters above the bars indicate the significant differences between treatments (small letters: among crosses occurring with a 0h interval; capital letters: among crosses occurring with a 24h interval. Error bars represent the standard errors of the mean. For offspring sex ratio, we obtained the estimates of the probability of being female and correspondent standard errors of the mean from the statistical GLM models. This takes into account sex ratio variation among females, as well as the quasi-binomial correction for overdispersion of the data. Numbers on the bottom of bars represent the sample size for each type of mating.

In *T. ludeni* females, the oviposition rate and the proportion of fertilized offspring varied significantly according to mating order in interaction with the mating interval ((F_2,248_ = 14.098, P < 0.0001 and F_2,152_ = 158.690, P < 0.0001, for oviposition rate and proportion of fertilized offspring respectively). Compared to the control treatment, *T. ludeni* females had lower oviposition rate when mating with *T. urticae* males immediately before conspecifics males (|t|(1) = 2.605, P = 0.0097; Fig. 3c). At the 24 hour interval, the conspecific crosses yielded higher oviposition rate than all other crosses in this time interval ((|t|(1) = 4.646, P < 0.0001 and |t|(1) = 3.805, P = 0.0002, in comparison with females mating with a conspecific before an heterospecific male and females mating with an heterospecific before mating with a conspecific, respectively; Fig. 3c). Additionally, when *T. ludeni* females mated with *T. urticae* males 24h after conspecific matings, the proportion of fertilized offspring was significantly lower than that of other crosses at this time interval (|t|(1) = 4.084, P < 0.0001 and |t|(1) = 3.586, P = 0.0005, in comparison with double conspecific matings and with females mating with a heterospecific before mating with a conspecific, respectively; Figure 3d). The mating sequence had no effect on the sex ratio at the 0h interval.

## Discussion

In this study, we investigated the consequences of mating with heterospecifics for the fertilization success and offspring viability in a system composed of three spider-mite species. We found that heterospecific matings between *T. urticae* and *T. ludeni* did not result in fertilized offspring (i.e., females), nor did it have any negative effects on egg viability, as shown for matings between *T. urticae* and *T. evansi* (Sato et al. 2014, Clemente et al. 2016). In fact, *T. ludeni* females that mate with *T. urticae* males produce more (male) offspring than virgin *T. ludeni* females. Second, the effects of heterospecific matings on the outcome of previous or subsequent matings with conspecifics were highly dependent on the species pair involved, on the trait measured and on the timing and order of mating events. Despite strong effects of particular mating sequences, the results taken as a whole suggest that the net effect of reproductive interactions between species is relatively weak.

Positive effects of interspecific reproductive interactions were found for fecundity. This can be due to a stimulation of oogenesis by the sperm of heterospecific males, increasing the availability of oocytes to subsequent matings with conspecifics. Indeed, oogenesis is stimulated by conspecific sperm in several species (Qazi et al. 2003, Xu and Wang 2011). This could also be the case with heterospecific sperm. If so, it could explain the higher fecundity found in crosses between *T. urticae* and *T. evansi*. In fact, earlier studies have documented that interactions with heterospecific males are not always negative. In some gynogenetic species, heterospecific mating is a prerequisite for embryogenesis (Gumm and Gabor 2005, Schlupp 2010). Moreover, in some invertebrate species, females receive nuptial gifts from heterospecific males (Vahed 1998, Costa-Schmidt and Machado 2012). However, to our knowledge, this is the first time that an increase in fecundity following a heterospecific mating is described in the literature. Such effects may thus be rare. Still, earlier studies may have overlooked them because they have not examined the roles of the order of mating in the outcome of heterospecific mating interactions.

Nonetheless, we also detected several negative effects of mating with heterospecifics, as found in most studies of reproductive interference (Gröning and Hochkirch 2008, Kishi 2015). We found both a reduction in the number of eggs laid and a decrease in fertilization success (i.e., offspring sex ratio). Whereas lower fecundity will most likely be costly in all ecological scenarios (assuming no trade-off with other traits), a decrease in fertilization success, leading to an excess of males in the population, will have an impact on fitness that is contingent upon the structure of spider-mite populations. Indeed, a higher frequency of males is likely to be more detrimental in recently-established populations, generally founded by a single female. In those populations, the optimal sex ratio is highly female-biased, hence a higher proportion of males is probably very costly. This is not the case in more panmitic populations, usually found when mites are established on a plant for a longer period (Roeder 1992, Macke et al. 2011). One also can speculate that a higher proportion of males may impose a strong cost in the other, competing species (assuming detrimental effects of heterospecific matings are stronger than positive effects). However, this behaviour is expected to be selected in males only if the cost they pay in terms of sperm loss is compensated by the benefit they would provide to their sisters. Hence, this behaviour is more likely to be selected in more structured populations.

Reproductive interference occurred independently of the order of matings and the time interval. This is at odds with expectations stemming from findings on conspecific matings, which show (a) first-male precedence and (b) exceptions to this rule only if the second male mates immediately after the first (Helle 1967). Therefore, the first male precedence found in conspecific matings cannot be extrapolated to matings involving heterospecific sperm. This contrasts with the recent finding that effects of heterospecific matings in *Drosophila* could be predicted from the harmful effects of conspecific mates (Yassin and David 2015), and that genes involved in conspecific male precedence also affect sperm precedence in multiple matings involving heterospecifics (Civetta and Finn 2014). Thus, the equivalence of effects of conspecific and heterospecific sperm on the outcome of conspecific matings is dependent on the type of effect and/or the species involved in the interaction. The mechanisms of sperm displacement in heterospecific matings in spider mites should therefore be investigated.

Since effects of heterospecific matings depend on the order and timing of occurrence, the outcome of interspecific reproductive interactions will depend on the frequency with which those different types of matings occur in nature. This, in turn, will depend on the discrimination abilities between species. First, these interactions will occur only if species discrimination is weak. This, indeed, has been explicitly demonstrated for the *T. evansi*/*T. urticae* interaction (Clemente et al. 2016), but not for *T. ludeni*/*T. urticae*. Still, these species do mate with heterospecifics under no choice scenarios, as shown here, hence the scope for the occurrence of reproductive interference does exist.

What then, would be the relative frequency of the mating sequences tested here? In spider mites, conspecific males often guard quiescent females (*i.e*., the last larval stage before becoming adult female), to ensure mating immediately after emergence. If males guard preferentially conspecific females, as shown in other spider mite species pairs (Collins et al. 1993, Takafuji et al. 1997), heterospecific matings will occur more often after rather than before conspecific ones. Hence, this leads to the prediction that the most common mating sequence among these species will be a heterospecific mating following a conspecific one. In this case, the detrimental effects of heterospecific matings for *T. urticae* females, are only visible when the two matings happen in close succession, while for *T. evansi* females, these effects are only apparent when the two matings occurred 24 hours apart. Since latency to copulation is not statistically different for these time intervals (Clemente et al. 2016), one expects these events to occur at similar frequency. However, since the effect of these heterospecific matings for *T. evansi* females is a reduction in fecundity, while for *T. urticae* it is an increase in the proportion of male offspring, the cost may be higher for *T. evansi* females (cf. above). Moreover, the excess of males produced by *T. urticae* can lead to more detrimental matings with *T. evansi* females, while reducing the probability of *T. urticae* females mating with a heterospecific male. This would mean that the invasive species would suffer more from reproductive interference than the resident.

Even assuming that all mating combinations do occur, reproductive interactions between *T. urticae* and *T. evansi* can be positive or negative for the two species, depending on the mating sequence. Moreover, reproductive interference between *T. urticae* and *T. ludeni* is stronger than that between *T. urticae* and *T. evansi*. Therefore, reproductive interference cannot be invoked to explain the exclusion of *T. urticae* in habitats with *T. evansi* (Ferragut et al. 2013, Sarmento et al. 2011b). Other factors may contribute to this exclusion, as the production of a dense web by *T. evansi*, which prevents heterospecifics from accessing the surface of the leaves to feed and oviposit (Sarmento et al. 2011b). Importantly, however, we show that the occurrence and strength of reproductive interference cannot be assessed with the unique evaluation of the outcome of a specific type of reproductive interaction. The different types of mating combinations – the order and interval between matings – have great influence on the overall outcome of heterospecific interactions and on the relative frequency of such events. This confirms the importance of using complete experimental designs on the detection and characterization of reproductive interference.

## Acknowledgements

The authors wish to thank Yukie Sato and Maurice W. Sabelis for helpful discussions during the early stages of this work. These experiments and Inês Santos were funded by Portuguese National Funds through an FCT-ANR project (FCT-ANR//BIA-EVF/0013/2012) to SM and Isabelle Olivieri. SHC and LRR had PhD fellowships funded by FCT (SFRH/BD/90156/2012 and SFRH/BD/87628/2012, respectively). SAMV and RP had Post-doctoral fellowships from FCT, (SFRH/BPD/66042/2009) and (SFRH/BPD/42801/2008) respectively. The authors declare no conflict of interest.

## Supporting Information

The following Supporting Information has been made available in the online version of this article.

Table S1 |Statistical tests and contrasts for the comparisons of fecundity and offspring sex ratio in crosses between conspecific and heterospecific males and females, using data from Block 1 or Blocks 1+2.

